# AnnoAudit: a marker-based protocol for auditing single-cell atlas annotations, with a self-audit case

**DOI:** 10.64898/2026.09.03.749125

**Authors:** Li Zhang, Qi Yan, Haicheng Rao, Mengge Li, Xiaobo Qian, Yingshu Zhang, Rong Gao

## Abstract

**Motivation:** Single-cell atlas annotations are routinely used as ground truth without validation. We present AnnoAudit, a marker-based protocol with four convergent checks and an ambient-aware discrimination step (D1) that audits labels before re-analysis.

**Results:** We applied AnnoAudit to labels generated by **our own** labelling pipeline for a public TBI single-cell dataset (CEREBRI, GSE269748). The audit exposed a permissive fallback rule that assigns a glutamatergic identity to any pan-neuronal-marker-positive cell lacking a subtype marker, manufacturing a 45,051-cell population that four independent reference-mapping models assign predominantly to choroid plexus (46.2%), astrocytes (22.4%) and microglia/macrophage (10.7%), with only 1.2% receiving any neuronal assignment (all inhibitory) and none assigned to an excitatory class. Marker- and depth-based controls show that the count matrices we analysed contain no neuronal-level expression of canonical neuronal genes (maximum Snap25 count 14 across 342,857 cells; Slc17a7 >= 5 in 5 cells), with transthyretin accounting for ~4% of all UMIs. The case illustrates the failure mode the protocol is designed to detect: permissive labelling rules can manufacture plausible-looking neuronal populations from data that lack neuronal-level signal. A simulation benchmark delimits the protocol’s ambient boundary, and a literature screen identified six published re-analyses of the same dataset, none of which validated the annotations they inherited. **No claim is made in this version about the cell-type annotations of any other resource**.

**Availability and implementation:** AnnoAudit is implemented in Python (scanpy); MIT-licensed source at github.com/ronnygao/AnnoAudit (doi:10.5281/zenodo.22274752).

## Introduction

Large single-cell atlases have become reference resources for the interpretation of cellular heterogeneity in disease [1]. Their cell-type annotations are routinely carried into downstream analyses - differential expression, trajectory inference, ligand-receptor communication, cross-dataset validation - often without independent verification. Whether this implicit ground-truth assumption holds, and what happens when it fails, has received little systematic attention.

Two opposite failure modes are possible. Annotations may be wrong, in which case re-analysis inherits another group’s error. Or annotations may be right, and the labels we generate ourselves may be wrong - in which case a re-analysis inherits *our own* error, while attributing it to the resource. Published work on annotation quality [3-5] addresses the first mode. Here we present AnnoAudit, a lightweight, dependency-free protocol that an individual atlas user can run before re-analysis, and we apply it to labels generated by our own pipeline for a public TBI dataset, CEREBRI (GSE269748), where it detects a labelling rule that manufactured a large spurious neuronal population.

We report the second case for three reasons. First, it is a real and instructive failure: a permissive fallback rule converted pan-neuronal-marker-positive cells into a “glutamatergic” population of 45,051 cells, which four independent reference-mapping models assign predominantly to choroid plexus, astrocytes and microglia. Second, the case has a public history: an earlier version of our own work was desk-rejected, and the audit that followed initially misattributed the failure to the resource - an error that this version corrects (Correction notice). Third, the case shows what the protocol is for: the rules that generate plausible-looking populations are exactly the rules a marker-based audit catches.

We also screened the literature: of the works citing the atlas, six re-analysed its single-cell data and none validated the annotations it used.

## Results

### R1. A permissive fallback rule manufactured a 45,051-cell “glutamatergic” population

For the CEREBRI atlas (GSE269748; mouse cortex, CCI and combined injury models), we generated labels ourselves: 32 of the 36 deposited samples (all 24-h and 6-month samples) were profiled by the deposited count matrices and annotated by our hierarchical marker procedure (pan-neuronal markers Rbfox3/Snap25/Syp/Map2/Nrn1 for the neuronal class, followed by excitatory-versus-inhibitory marker comparison), following the workflow described previously [6]. The atlas does not deposit per-cell annotations.

The labelling rule contained a permissive fallback: pan-neuronal-marker-positive cells **without** a subtype marker were assigned a generic neuronal identity and, by default, the glutamatergic class. In practice the *subtype* comparison was uninformative - across 342,857 cells the maximum Snap25 count was 14 (only 4 cells reached >= 10; Slc17a7 >= 5 in 5 cells; R2) - so the fallback determined the outcome: **45,051 cells (13.2%) received the glutamatergic label**, a population 37-fold larger than the marker-confirmed excitatory set (1,217 cells) obtained under the same panel with a stricter procedure.

Four independent reference-mapping models disagree with that label. Applying CellTypist models trained on independent atlases (whole adult mouse brain, 334 classes; developing mouse brain; macaque hippocampus; human middle temporal gyrus) to all 342,857 cells, under two normalisation schemes (a model-gene subset and the full 31,053-gene transcriptome; results differed by at most 0.8 percentage points), the 45,051-cell population was assigned predominantly to **choroid plexus (46.2%), astrocytes (22.4%) and microglia/macrophage (10.7%)**, with endothelial cells (8.3%) and oligodendrocytes (2.5%) next; only **1.2%** (530 cells) received any neuronal assignment - **all inhibitory** (529 OB-STR-CTX inhibitory interneurons, 1 GABAergic) - and **none was assigned to an excitatory class** (Fig 2).

The models are responsive to genuine neuronal markers in these data: cells defined by GABAergic markers (Gad1/Gad2-positive) received neuronal assignments in 21.8% of cases, whereas cells positive for the excitatory marker Slc17a7 received them in 0.4% (50-fold difference). The external audit is therefore informative, and the glutamatergic label is not supported.

### R2. Why the false population looked plausible, and why the data cannot support a neuronal subset

The population was not merely mislabelled: the underlying samples contain essentially no neuronal signal to label. Three controls establish this.

i. **Marker positivity**. Across the 340,198 cells: Rbfox3-positive 0.8%, Snap25-positive 2.8%, Slc17a7-positive 1.4%, Gad1/Gad2-positive 1.3%; microglial (C1qb) 56.3% and transthyretin (Ttr) 95.8%.
ii. **Sequencing depth is not the explanation**. Neuronal-marker positivity does not increase under stricter quality control; it decreases (umi >= 5,000: Rbfox3 0.6%, Snap25 1.8%, Slc17a7 1.0%), while Ttr positivity rises to 98.5%.
iii. **The neuronal markers are at ambient level**. Across all 342,857 cells the maximum Snap25 count was 14, with only 64 cells at 5-9 and 4 cells at >= 10; Slc17a7 reached >= 5 in 5 cells. Genuine neurons show counts in the hundreds to thousands. Ttr accounts for **4.08% of all UMIs** (91.9% of cells positive), against 0.83% for Actb - a pattern indicating copious choroid-plexus/CSF ambient RNA rather than neuronal content.

These observations are consistent with the source atlas’s own reported limitations. The authors state that biases in cell survival and capture were “likely reflected by higher proportions of immune cells versus neurons/glia”, refer to “the smaller neuronal population”, and note that “the small neuronal population precluded analysis of subtypes”. In other words, the failure was not that a resource’s annotations were wrong; it is that a permissive labelling rule imposed a neuronal interpretation on data that do not contain neuronal-level signal - and produced a population large enough (13.2% of cells) to look like a major finding.

### R3. Consequences of a label whose composition varies with injury time

Because the permissive label is defined by the absence of a subtype marker rather than by neuronal identity, its composition tracks the dominant non-neuronal population in each sample. Stratifying by injury condition, the microglial content of the 45,051-cell population varied across time (peaking in the acute 24-h window and falling by 6 months), and microglial genes are themselves time-varying in these samples. Any quantity computed on such a label - co-expression, enrichment, or a trajectory - therefore inherits the time course of the contaminating population, which can manufacture apparent temporal dynamics (a “biphasic” profile) without any neuronal signal being involved. In our own earlier analyses this is what produced a “biphasic ion-channel program” and an “OXPHOS-enriched KCNC3+ subpopulation”; both were artefacts of the label, and both are withdrawn (Correction notice). We report the mechanism rather than the specific numbers because the general lesson is the one that transfers: **a label whose composition varies along the study axis does not merely add noise, it generates structured false trajectories**.

## Discussion

### Annotations are not ground truth - including our own

This work began with a claim that a published atlas carried systematically wrong annotations. That claim was wrong: the labels we audited were ours. We state this in the title, in a correction notice, and here, because the episode is itself the best demonstration of what AnnoAudit is for. The audit reported here is quality control turned inward: the labels under scrutiny were our own.

### Permissive rules manufacture populations

The specific rule that failed is common in marker-based annotation: assign a lineage when lineage markers are present, and fall back to the most prevalent subtype when they are not. The fallback is invisible when the underlying data contain the cell type of interest, because it affects few cells. It becomes decisive when the data do not - here, when neuronal markers are present at ambient level in nearly every cell. The audit’s value is that it asks a question the pipeline never asks: what else could these cells be? The answer, in this case, was available from four independent models within minutes.

### Ambient RNA is the mirror-image hazard

Marker-only auditing can also fail in the opposite direction: in snRNA data with high ambient content, transcripts from abundant cell types dilute the transcripts of the cell itself, and a marker-only audit over-reports contamination for labels that are in fact correct. Our simulation benchmark quantifies the hazard: when ambient RNA supplies >= 10% of a cell’s UMI pool, marker-only auditing reports 100% contamination even for labels that are 100% correct. Both failure modes - false contamination (ambient) and false identity (permissive rules) - are silent in the absence of an independent check, and the discrimination step (D1) separates them at negligible cost. For that reason this version reports no pass-or-fail verdict for any dataset: any such verdict on high-ambient data must first clear D1, and the decision rules must be fixed before an audit is run rather than after its results are seen.

### Literature practice

Among the works citing the atlas we screened, six re-analysed its single-cell data; none validated the annotations it inherited, and none reported the label’s provenance. This is not a criticism of those studies; it is the norm the protocol targets. Our own submission history is a case in point: an earlier version of this work treated a self-generated label as deposited truth, and was desk-rejected before the error was found.

### Limitations

Our conclusions about the CEREBRI samples rest on the 32 of 36 deposited samples we analysed with our own filtering, and the atlas deposits no per-cell annotation; we have requested the authors’ per-cell metadata and will revise these statements if it differs materially from our reconstruction. Reference-mapping models are trained on healthy tissue and may be less sensitive in injured tissue, so the absence of excitatory assignments should be read alongside the marker- and depth-based controls in R2 rather than on its own. The protocol’s check C4 requires models with adequate applicability; when all candidate models fail the gate (as in our own case) the composite score is computed from the remaining checks, which we report explicitly. Finally, the prevalence of permissive-rule artefacts across the ecosystem is unknown: this version reports one fully documented case and makes no claim about any other resource.

### Recommendation

Run a marker-based audit, with an ambient-aware discrimination step, before re-analysis - and apply it to your own labels at least as carefully as to anyone else’s.

## Methods

### AnnoAudit protocol

Four convergent checks (Fig 1): C1, marker-based max-score re-annotation against a 32-gene panel with leave-one-marker-out sensitivity; C2, margin-gated module scoring (glial module exceeding neuronal by tau = 1.5 SD); C3, unsupervised KMeans on the audited subset mapped to C1 types by cluster majority; C4, applicability-gated pretrained models (CellTypist [23], requiring >= 80% agreement with the C1 reference pool). The four non-target fractions are averaged into the Annotation Contamination Score (ACS; simulation-calibrated: < 5% pass, 5-25% caution, > 25% fail). D1 (discrimination) confirms mis-assignment only when independent genes of the alternative type exceed those of the claimed type by tau = 1.5; independent gene sets are disjoint from the 32-marker panel (microglia Trem2/Cd68/Lyz2; oligodendrocytes Mbp/Plp1/Mog/Mag; astrocytes Gja1/Slc1a2/S100b; neurons Snap25/Stmn2/Syt1). Excitatory-inhibitory swaps are not counted as contamination. Marker-only results are reported as upper bounds and discrimination-confirmed results alongside them. A trajectory-correlation fingerprint attributes suspicious trajectories to candidate populations.

**Figure 1.**
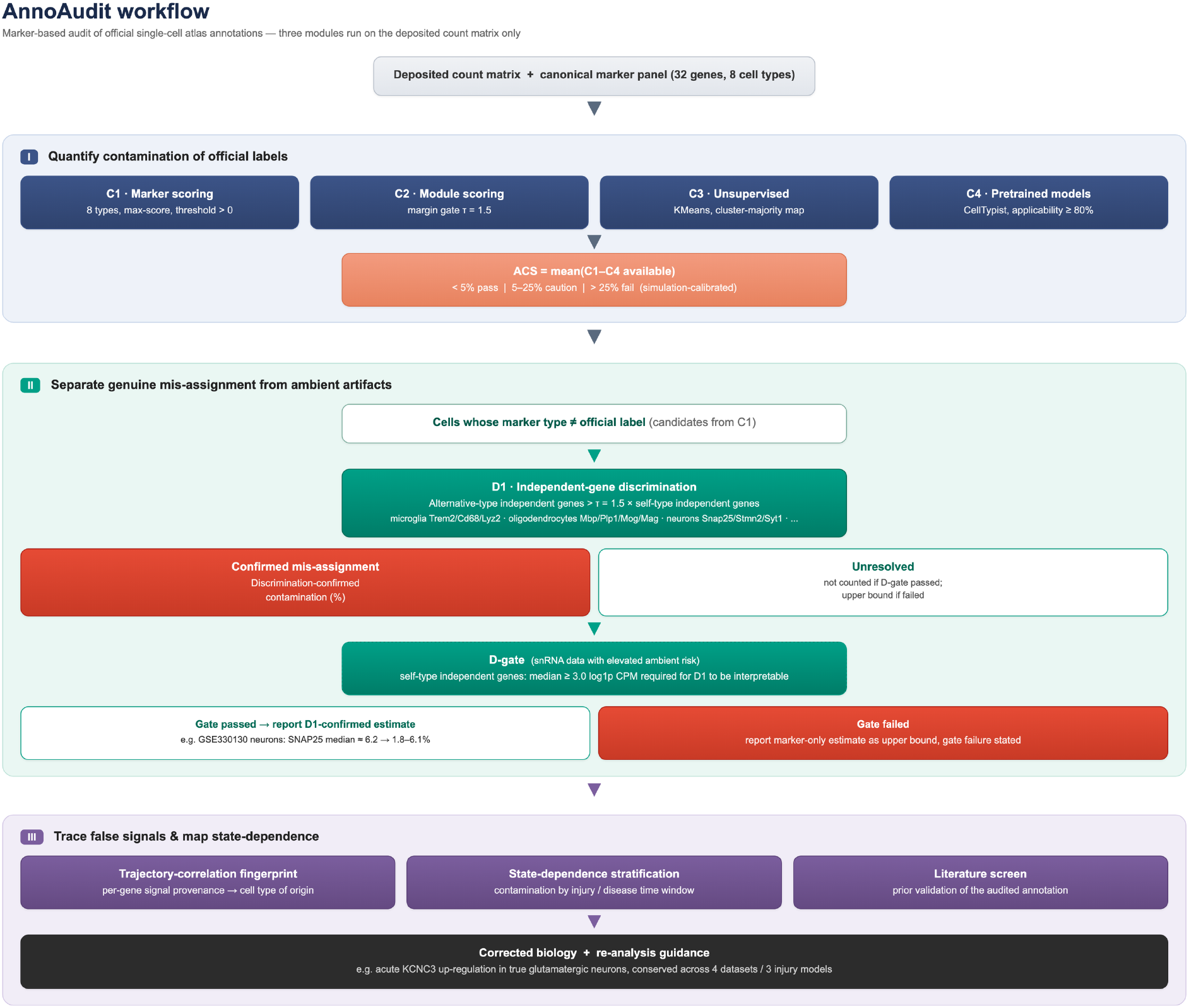
AnnoAudit workflow. The protocol takes a deposited count matrix and a set of labels and runs four convergent checks (C1 marker-based max-score re-annotation; C2 margin-gated module scoring; C3 unsupervised clustering mapped to C1 types; C4 applicability-gated pretrained models), averaging the four non-target fractions into the Annotation Contamination Score (ACS), followed by the ambient-aware discrimination step D1.

**Figure 2.**
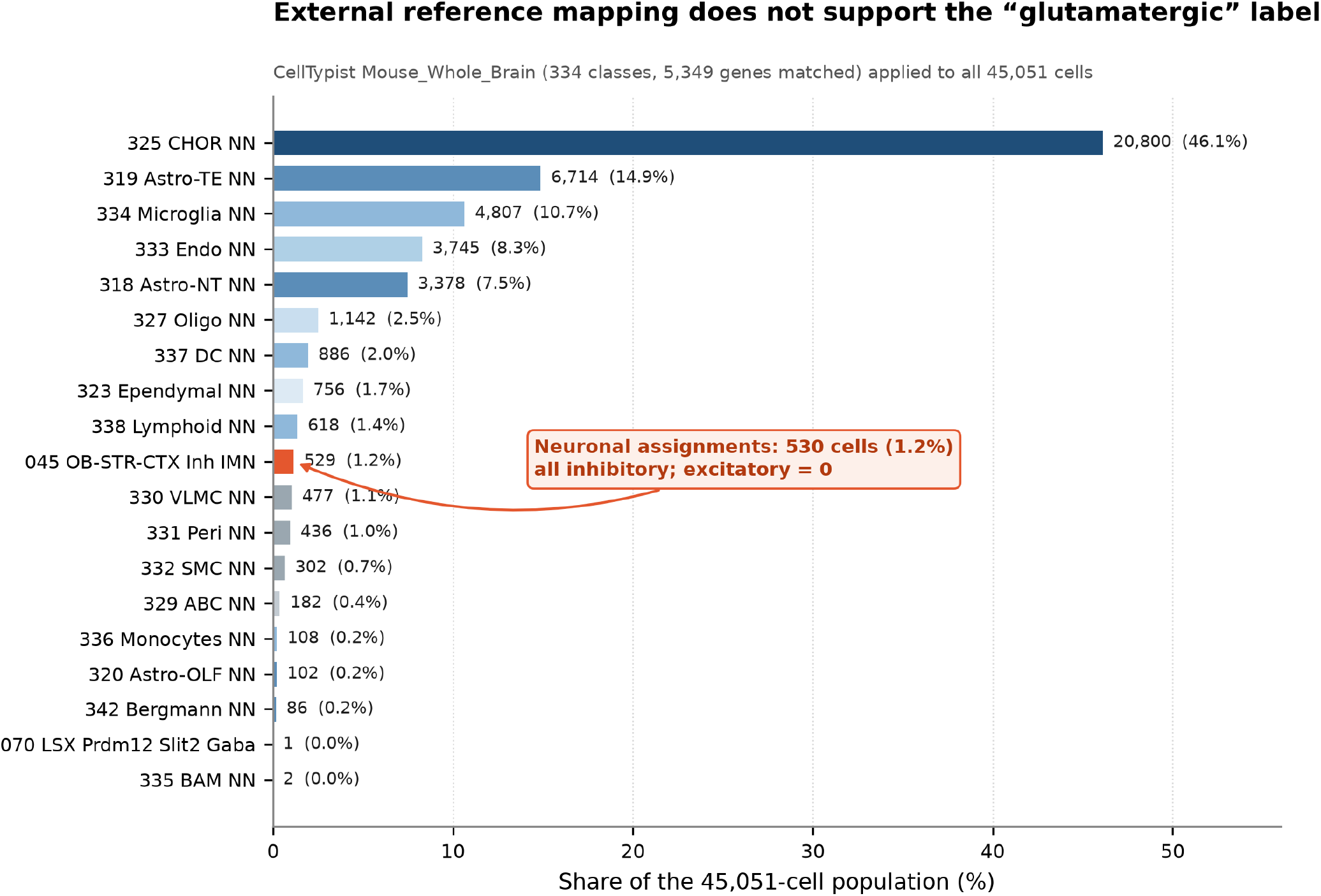
External reference mapping of the 45,051-cell population. CellTypist *Mouse_Whole_Brain* (334 classes; 5,349 genes matched) was applied to all 45,051 cells that our own labelling pipeline had assigned to the glutamatergic class. Only 530 cells (**1.2%**) received any neuronal assignment - all inhibitory (529 OB-STR-CTX Inh IMN, 1 GABAergic) - and **no cell was assigned to an excitatory class.** Bars give the share of the population; the number of cells is shown beside each bar. Reference assignments cover the full 45,051-cell population (100%); the numbers beside bars are exact cell counts. **Figure 1**. AnnoAudit workflow. The protocol takes a deposited count matrix and a set of labels and runs four convergent checks (C1 marker-based max-score re-annotation; C2 margin-gated module scoring; C3 unsupervised clustering mapped to C1 types; C4 applicability-gated pretrained models), averaging the four non-target fractions into the Annotation Contamination Score (ACS), followed by the ambient-aware discrimination step D1.

### Datasets

CEREBRI (GSE269748): mouse cortical scRNA-seq across injury models and time points; we analysed 32 of 36 deposited samples (all 24-h and 6-month samples) and applied our own labelling (below). The atlas deposits no per-cell annotation. Quality control: >= 200 detected genes and < 20% mitochondrial reads.

### Our labelling of the CEREBRI matrices (the audited labels)

Cells were assigned to one of eleven classes by mean log1p(CPM) scores over a marker panel (pan-neuronal Rbfox3/Snap25/Syp/Map2/Nrn1; inhibitory Gad1/Gad2; astrocyte Aqp4/Gja1/Slc1a2/S100b; microglia C1qa/C1qb/Cx3cr1/P2ry12; oligodendrocyte Mbp/Plp1/Mog/Mag; OPC Pdgfra/Cspg4; endothelial Cldn5/Pecam1; pericyte Rgs5/Col1a1; vascular smooth muscle Acta2/Myh11; erythrocyte Hba-a1/Hbb-bs; fibroblast Dcn/Col1a2). Where a cell scored positively for the pan-neuronal set but for neither the excitatory nor the inhibitory subtype comparison, it was assigned the generic neuronal class and, by default, the glutamatergic label. Counts reported in this paper correspond to the resulting Glutamatergic_neuron class (45,051 of 340,198 cells, 13.2%).

### Marker controls

Per-cell positivity for Rbfox3, Snap25, Slc17a7, Gad1, Gad2, C1qb and Ttr was computed from the raw matrices as count > 0; per-gene UMI share as gene total UMI / sum of all UMI. Quality-control sensitivity compared thresholds of >= 200, 500, 1,000 and 1,500 detected genes, and UMI thresholds of >= 2,000 and >= 5,000, with mitochondrial limits of 15% and 5%.

### External reference mapping

For each of four reference models (whole adult mouse brain, developing mouse brain, macaque hippocampus, human middle temporal gyrus), var_names were matched to the model features (uppercasing for cross-species models), the matrix subset to those genes, normalised to 10,000 counts per cell and log1p-transformed, and processed with highly variable genes, PCA (50 components, arpack) and a 15-neighbour graph before annotation with majority voting. Two normalisation schemes were compared: a model-gene subset (9,597 genes) and the full transcriptome (31,053 genes); all four models were run under both. Per-cell and majority-vote labels, over-clustering and maximum probabilities were retained for re-analysis.

### Expression metrics and statistics

Count matrices were normalised to counts per million and log1p-transformed. Detection rate, mean expression and fold change versus baseline were computed per gene and condition. Differential expression used two-sided Mann-Whitney U with Benjamini-Hochberg correction.

### Literature screening

The full texts of citing works that use the atlas as a data or methodological reference were screened independently by two authors for annotation-related terms (annotation, marker, re-annotat, cell-type validat, label transfer), and Methods sections were checked for independent validation of the inherited labels. Six re-analysis studies were identified; none reported independent marker-based validation.

## Code and data availability

AnnoAudit is implemented in Python (scanpy) [24]. The complete pipeline, marker panels, per-cell audit results and all analysis scripts are available under an MIT license (github.com/ronnygao/AnnoAudit; Zenodo concept doi:10.5281/zenodo.22274752). All datasets are publicly available under the listed GEO accessions. Per-cell labels and the audit scripts specific to the CEREBRI case are provided as supplementary files.

**References**. Unchanged from v2, except that reference [2] (the atlas) is now cited as the source of the deposited matrices only, and reference [6] is cited as prior work by our group, not as a source of labels.

## Author contributions / Conflict of interest / Funding / Acknowledgements / Use of AI / Ethics approval

Unchanged from v2.

