## Supplementary Tables S1-S4 for "AnnoAudit: a marker-based protocol for auditing single-cell atlas annotations, with a self-audit case" (from another version of this preprint: version 1, 2026-09-08)

**Supplementary Tables (AnnoAudit)**

**Supplementary Table S3. Sensitivity of the C2 margin gate and composite ACS to the gate threshold τ**

τ is the margin gate of check C2: a cell is counted as non-neuronal when its best glial module score exceeds its best neuronal module score by more than τ standard deviations (`scanpy.tl.score_genes`). C1 and C3 do not depend on τ (CEREBRI: C1 = 97.6%, C3 = 100.0%; GSE330130: C1 = 73.3%, C3 = 100.0%); ACS = mean(C1, C2, C3). C4 was not computable (all CellTypist models failed the applicability gate). Across the full τ = 1.0–2.0 range, both audited atlases remain far above the ACS > 25% failure threshold, and all qualitative conclusions are unchanged at every tested τ. Reproducible with `scripts/v15_tau_sensitivity.py`.

| Dataset (official subset, n) | Check | τ = 1.0 | τ = 1.5 | τ = 2.0 |
| --- | --- | --- | --- | --- |
| CEREBRI Glutamatergic (45,051) | C2 (%) | 60.2 | 50.3 | 43.1 |
| CEREBRI Glutamatergic (45,051) | ACS (%) | 85.9 | 82.6 | 80.2 |
| GSE330130 Neuron (2,450) | C2 (%) | 47.1 | 37.8 | 31.3 |
| GSE330130 Neuron (2,450) | ACS (%) | 73.5 | 70.4 | 68.2 |

*τ = 1.5 is the baseline used throughout the manuscript (default of `scanpy.tl.score_genes`).*

**Supplementary Table S1. Contingency table of official CEREBRI annotation versus unsupervised clustering labels (120,000 cells)**

Official cell-type labels (rows) versus KMeans cluster assignment (columns, no marker information used). The distribution of official glutamatergic cells across nine clusters — 4,982 assigned to the microglial cluster, 2,975 to the astrocytic cluster, 337 to the excitatory cluster — demonstrates that the misassignment is structural (annotation composition error), not a systematic metadata/label offset (which would produce a near-diagonal correspondence).

| Official label | Astrocyte | Endothelial | Excitatory | Inhibitory | Microglia | OPC | Oligodendrocyte | Pericyte | Unknown |
| --- | --- | --- | --- | --- | --- | --- | --- | --- | --- |
| Astrocyte | 3046 | 585 | 19 | 17 | 6531 | 264 | 1670 | 682 | 2259 |
| Endothelial | 5 | 5424 | 1 | 0 | 185 | 4 | 44 | 396 | 201 |
| Erythrocyte | 38 | 28 | 11 | 1 | 198 | 25 | 113 | 119 | 297 |
| Fibroblast | 85 | 54 | 10 | 1 | 344 | 116 | 206 | 249 | 543 |
| GABAergic_neuron | 148 | 59 | 7 | 209 | 225 | 9 | 120 | 31 | 16 |
| **Glutamatergic_neuron** | **2975** | **1480** | **337** | **40** | **4982** | **597** | **2493** | **1785** | **1260** |
| Microglia | 82 | 948 | 13 | 5 | 64750 | 54 | 333 | 550 | 513 |
| OPC | 2 | 30 | 0 | 6 | 774 | 320 | 30 | 23 | 50 |
| Oligodendrocyte | 2 | 7 | 1 | 0 | 137 | 6 | 1680 | 10 | 28 |
| Pericyte | 0 | 45 | 1 | 0 | 35 | 12 | 6 | 658 | 17 |
| Unknown | 0 | 0 | 11 | 11 | 2274 | 2 | 0 | 0 | 3452 |
| VSM | 15 | 178 | 3 | 1 | 722 | 63 | 83 | 1440 | 73 |

**Supplementary Table S2. Sample-level (pseudobulk) KCNC3 validation in GSE277487**

Per-animal mean KCNC3 expression in marker-defined excitatory neurons. Sample-level analysis confirms the direction of the cell-level effect but lacks statistical power given the small number of control animals with passing QC (n = 2).

| Group | Sample | Excitatory cells (n) | Mean KCNC3 (log1p CPM) | Detection rate (%) |
| --- | --- | --- | --- | --- |
| Control | GSM8523266 | 7586 | 0.2966 | 23.56 |
| Control | GSM8523267 | 9522 | 0.3966 | 29.88 |
| TBI 6 h | GSM8523257 | 10116 | 0.6916 | 43.71 |
| TBI 6 h | GSM8523258 | 8812 | 0.5337 | 36.88 |
| TBI 6 h | GSM8523259 | 8428 | 0.4696 | 34.67 |
| TBI 2 d | GSM8523260 | 9997 | 0.3267 | 23.97 |
| TBI 2 d | GSM8523261 | 6432 | 0.5174 | 34.50 |
| TBI 2 d | GSM8523262 | 8793 | 0.5792 | 37.70 |
| TBI 4 d | GSM8523264 | 10110 | 0.4315 | 30.71 |

**Sample-level comparisons (two-sided Mann–Whitney U):**

- Control (n = 2, mean 0.347) vs TBI 6 h (n = 3, mean 0.565): fold change +1.63×, p = 0.20
- Control (n = 2, mean 0.347) vs TBI 2 d (n = 3, mean 0.474): fold change +1.37×, p = 0.40
- Control (n = 2, mean 0.347) vs acute pooled (6 h + 2 d, n = 6, mean 0.520): fold change +1.50×, p = 0.14

*Note: 9 of 12 animals passed QC in the deposited annotation; Control n = 2 limits statistical power. The primary evidence for acute KCNC3 up-regulation rests on consistent replication across four datasets (GSE277487, GSE160763, GSE247339, CEREBRI true population) and three injury models.*

**Supplementary Table S4. Discriminative performance of the C2 margin gate and cross-species transferability of the marker panel**

| Metric | CEREBRI (mouse TBI cortex; n = 45,051) | GSE330130 (human ALS spinal cord; n = 2,450) |
| --- | --- | --- |
| AUROC of continuous margin vs C1 marker-typing labels | 0.996 (95% CI 0.995–0.997) | 1.000 (95% CI 1.000–1.000) |
| Permutation test (1,000 label permutations; null AUROC) | P < 0.001 (null 0.500 ± 0.009) | P < 0.001 (null 0.500 ± 0.013) |
| Sensitivity at τ = 1.5 (vs C1 labels) | 56.0% | 57.4% |
| Specificity at τ = 1.5 (vs C1 labels) | 100% | 100% |
| Balanced accuracy at τ = 1.5 | 78.0% | 78.7% |
| Independent 16-gene panel: AUROC | 0.753 | 0.721 |
| Independent 16-gene panel: Pearson r (margin) | 0.49 | 0.54 |
| Independent 16-gene panel: sign agreement with C2 margin | 79.2% | 55.6% |
| C2 glial fraction by official sub-label | — | Motor 60.7% > Inhibitory 37.1% > Excitatory 23.2% |
| C3 cluster structure (C2 glial fraction per cluster) | 4/10 clusters ≥80%; 2/10 clusters <17% | — |
| Independent panel marker coverage | 16/16 | 16/16 |
| 32-gene panel coverage | 32/32 | 32/32 |
| 1:1 human orthologs detected in both datasets | 32/32 | 32/32 |

*The continuous margin is the difference between the best glial and best neuronal module scores (C2, Methods). The independent anchor panel comprises five neuronal (NRGN, SNAP25, STMN2, SYT1, GRIN1) and eleven glial markers (S100B, GLUL, GJA1, ITGAM, CTSS, LGALS3, CNP, OLIG2, SOX10, VWF, KCNJ8), none of which overlap the 32-gene audit panel. GSE330130 official sub-labels: motor neurons carry the highest C2 glial fraction. Reproducible with `scripts/v16_methodology_enhancement.py`.*
