## Supplementary material for "AnnoAudit: a marker-based protocol for auditing single-cell atlas annotations, with a self-audit case": Tables 1-4 (from another version of this preprint: version 1, 2026-09-08)

**Table 1'. Annotation Contamination Score (ACS) across audited official labels (final protocol)**

| Atlas | Official label | N (official) | C1 marker | C2 module (margin τ=1.5) | C3 KMeans | C4 CellTypist (applicability ≥80%) | ACS | Verdict |
| --- | --- | --- | --- | --- | --- | --- | --- | --- |
| CEREBRI (GSE269748) | "Glutamatergic_neuron" | 45,051 | 97.6% | 50.3% | 100.0% | N/A (DMB 12.4% / MTG 18.1% / PFC 1.6% all FAIL) | **82.6%** (mean of C1/C2/C3) | ❌ FAIL (>25%) |
| GSE330130 (human ALS cord) | "Neuron" | 2,450 | 73.3% | 37.8% | 100.0% | N/A (MTG 78.8% / PFC 30.9% all FAIL) | **70.4%** (mean of C1/C2/C3) | ❌ FAIL (>25%) |
| GSE277487 (own marker annotation) | excitatory (no official label) | 162,892 | <5% | — | — | — | **≤5%** | ✅ PASS (control) |
| GSE160763 (own marker annotation) | microglial/excitatory (no official label) | 20,642 | <5% | — | — | — | **≤5%** | ✅ PASS (control) |
| GSE247339 (own marker annotation) | excitatory (no official label) | 85,431 | <5% | — | — | — | **≤5%** | ✅ PASS (control) |

**Simulation calibration** (n = 5 replicates per level; ACS values in percent; C4 omitted because all audited models fail the applicability gate):

| True contamination c | ACS (mean ± SD) | C1 | C2 (margin) | C3 (KMeans) |
| --- | --- | --- | --- | --- |
| 0% | 0.3 ± 0.1 | 0.0 | 1.0 | 0.0 |
| 5% | 4.7 ± 0.3 | 5.0 | 5.5 | 3.8 |
| 10% | 8.8 ± 0.3 | 10.0 | 8.9 | 7.4 |
| 20% | 17.5 ± 0.8 | 20.0 | 17.0 | 15.5 |
| 50% | 44.5 ± 1.4 | 50.0 | 36.9 | 46.6 |
| 80% | 73.9 ± 1.5 | 80.0 | 57.0 | 84.6 |

**Key points**: (i) the two deposited official neuronal labels audited (CEREBRI glutamatergic, GSE330130 neuron) both FAIL with composite ACS 70–83%; (ii) marker-defined controls in three further datasets show biologically plausible proportions consistent with the original studies; (iii) ACS ≈ 0.9 × c in the 0–20% range (r² = 0.999), supporting the >25% failure threshold for contamination above ~15%; (iv) the C2 margin gate underestimates contamination at extreme high levels (57% at c = 80%) because it counts a cell as non-neuronal only when the best glial module exceeds the best neuronal module by τ, so ACS is the more accurate estimator than C2 alone.

──────────────────────────────

**Table 1. Marker audit of official annotations across atlases**

| Atlas | Species / tissue | Official annotation | Official N | After marker validation | Retention | Contamination composition | Verdict |
| --- | --- | --- | --- | --- | --- | --- | --- |
| CEREBRI (GSE269748) | Mouse cortex (CCI etc.) | "Glutamatergic_neuron" | 45,051 | 1,174 true excitatory | 2.1% (963 overlap with the 1,174 marker-confirmed) | Microglia 31.4% / Astrocyte 18.6% / Oligodendrocyte 15.9% / Pericyte 11.2% / Endothelial 9.2% / Unknown 7.8% / OPC 3.6% / Inhibitory 0.3% | ❌ **Systematic contamination** (97.9% non-glutamatergic) |
| GSE330130 (human ALS lumbar cord) | Human spinal cord (ALS) | "Neuron" (excitatory + inhibitory + motor) | 2,450 | 654 retained as neurons (311 excitatory / 343 inhibitory) | 26.7% | 73.3% reclassified as non-neuronal (1,796 = oligodendrocyte 1,360 + astrocyte 253 + endothelial 74 + OPC 44 + pericyte 34 + microglia 22; 9 unclassified) | ❌ Heavily contaminated (>two-thirds of neuron labels incorrect) |
| GSE277487 (CCI snRNA) | Mouse cortex | Data-driven marker annotation | 162,892 (post-QC) | 91,739 excitatory (56.3%) | — | Excitatory-dominated, consistent with the original study | ✅ Reasonable control |
| GSE160763 (FPI scRNA) | Mouse cortex | Data-driven marker annotation | 20,642 (post-QC) | 940 excitatory (4.6%) | — | Microglia-dominated (~54%, consistent with the original study) | ✅ Reasonable control |

**Key point**: atlases distributed with official annotations (CEREBRI, GSE330130) show systematic neuronal-label contamination; data-driven annotations (GSE277487, GSE160763) are biologically plausible — "has an official annotation" ≠ "annotation is trustworthy"; auditing is a necessary step.

──────────────────────────────

**Table 2. The 40 false biphasic ion channels manufactured by the official annotation → 0 under marker-validated annotation**

**Criteria**: acute up-regulation (24 h: FDR < 0.05 and |log2FC| ≥ 0.25; primary criterion) + chronic down-regulation (6 mo: same criteria) — the "compensation-to-decompensation" pattern.

| Family | Official annotation (40) | Marker true annotation |
| --- | --- | --- |
| *KCTD* potassium-channel auxiliary (×10) | *Kctd1*, *Kctd2*, *Kctd6*, *Kctd7*, *Kctd9*, *Kctd10*, *Kctd11*, *Kctd15*, *Kctd20*, *Kctd21* | **0** |
| Voltage-gated potassium (×3) | *Kcnh1*, *Kcnh2*, *Kcnc3* (Kv3.3) | 0 |
| Calcium-activated potassium (×1) | *Kcnn1* (SK) | 0 |
| TRP channels (×6) | *Trpm3*, *Trpm4*, *Trpm7*, *Trpv4*, *Trpc1*, *Trpc4ap* | 0 |
| Chloride channels (×6) | *Ano6*, *Ano8*, *Ano10*, *Clcn4*, *Clcn5*, *Clcn7* | 0 |
| Nicotinic acetylcholine receptors (×3) | *Chrna4*, *Chrnb4*, *Chrna1os* | 0 |
| Two-pore/TRIC (×4) | *Tpcn1*, *Tpcn2*, *Tmem38a*, *Tmem38b* | 0 |
| Voltage-gated calcium auxiliary (×3) | *Cacna2d2*, *Cacna2d3*, *Cacng7* | 0 |
| Other (×4) | *P2rx6*, *Mcoln1*, *Scnn1a*, *Gabra4* | 0 |
| **Total** | **40 (13.0% of 307)** | **0** |

*Note:* Chrna1os *is an antisense transcript in the same cluster as* Chrnb4*;* Tmem38b *is counted once (Two-pore/TRIC family).*

**False signals that disappear together**:

- *KCNC3*+ OXPHOS enrichment: official FDR < 0.001 (n = 2,416) → true annotation **p = 0.11** (n = 43), fold 1.24 → 1.06
- *KCNC3* "7-day down-regulation −46%": true direction is **up-regulation** (see Table 3)

──────────────────────────────

**Table 3. Cross-dataset validation of *KCNC3* in true excitatory neurons (marker annotation, all measured)**

| Dataset | Model | Time point | n cells | mean (log1p CPM) | vs control | Detection rate | vs control |
| --- | --- | --- | --- | --- | --- | --- | --- |
| GSE277487 | CCI severe snRNA | control | 27,700 | 0.328 | — | 21.5% | — |
|  |  | **6 h** | 23,016 | **0.688** | **+110%** | **40.8%** | +90% |
|  |  | 2–4 d | 41,023 | 0.458 | +40% | 33.7% | +57% |
| GSE160763 | FPI mild scRNA | control | 361 | 0.158 | — | 7.5% | — |
|  |  | **7 d** | 579 | **0.213** | **+35%** | 10.9% | +45% |
| GSE247339 | mTBI mild scRNA | sham | 802 | 0.128 | — | 8.9% | — |
|  |  | **24 h** | 983 | **0.198** | **+55%** | 11.0% | +24% |
|  |  | **7 d** | 757 | 0.196 | +53% | 8.1% | −9% |
|  |  | **21 d** | 753 | **0.215** | **+68%** | 13.7% | +55% |
| CEREBRI (official annotation, control) | CCI scRNA | 24h contra/7 d/6 mo | — | +94%/**−46%**/−45% | — | — | Official trajectory (**erroneous**); 6 h +43% from GSE277487 official annotation |

**Key point**: consistent up-regulation across three datasets and three injury models (+35% to +110%), opposite in direction to the official annotation's claim of 7-day down-regulation (−46%); subacute (2–7 d) expression remains above control. *Kcnc1* is co-up-regulated (6 h +96%, GSE277487).

──────────────────────────────

**Table 4. Propagation of the CEREBRI annotation in the re-analysis literature (as of August 2026)**

| Metric | Value |
| --- | --- |
| Total citations of CEREBRI (CrossRef) | 73 |
| Citing works using the atlas as a data or methodological reference | 65 |
| Re-analysis studies explicitly requesting cell-type subsets | 6 |
| Studies validating the official annotations against independent markers | **0** |

**Key point**: despite heavy reuse, no published re-analysis study has independently validated the official annotations — the contamination propagated silently through the citation record.

*To our knowledge; screening methods are described in the main text (Literature screening for annotation validation).
